# Quantum transduction from electron spin state to signaling in wild-type AsLOV2

**DOI:** 10.64898/2026.06.30.735402

**Authors:** William Salvia, Joshua Straub, Shiny Maity, Alexey Bogdanov, Subhajyoti Chaudhuri, Ruixian Han, Songi Han

## Abstract

Electron spin coherence is often thought to be at odds with the warm, wet conditions of biological systems^1^. Coherent electron-spin dynamics have been shown to modulate fluorescence in some flavoproteins, but not the chemistry by which a protein signals^2,3,4^. The direct connection between electron spin coherence and biological function is therefore still missing. Here we show that a magnetic field modulates formation of the covalent FMN–cysteine photoadduct that constitutes the signaling state of wild-type *Avena sativa* LOV2 (AsLOV2), via a spin-selective radical-pair reaction. AsLOV2 was thought magnetically inactive^5^; instead a fluorescence magnetic field effect emerges near 75 mT. A 650 mT field raises the signaling state population and shortens product formation by 6.7 ± 0.6%.The short FMN–cysteine separation produces a large dipolar coupling that blocks hyperfine-driven triplet-singlet interconversion at low field; the unusually large flavin-cysteinyl radical *g*-factor difference overcomes it at high field. Bond formation proceeds from the singlet, so the coherent interconversion that the field controls sets the rate of signaling-state formation. This arrangement is canonical in LOV domains^6^, so the magnetic control of this signaling chemistry likely extends across natural and engineered photoreceptors.

## 1 Introduction

A growing body of evidence shows that quantum properties can affect biological function, yet the findings are contested, the mechanisms are disputed, and conclusive cases are few^1,7,8^. Open questions include whether such effects are rare or common, and whether quantum coherence can directly modulate protein signaling that underlies biological processes. Coherent spin dynamics has been shown to influence flavin fluorescence in living bacteria transformed and overexpressed with certain engineered variants of a Light-Oxygen-Voltage (LOV) flavoprotein^2,3^ and in transgenic animals^9^. Radio waves have been used to modulate photoluminescence in cryptochromes^3^ and in the engineered MagLOV^2,5^ and iLOV^3^ variants; in each case the readout is emission, and the radical pair being controlled does not directly lie on a signal transduction pathway. Spin state-or coherence-dependent control on biological signal transduction has never been demonstrated.

The central challenge for coherent quantum effects in biology is fast decoherence in warm and crowded solution conditions^1,7^. Electron spin coherence is an exception. In a spin-correlated radical pair (SCRP) recombination is spin-selective: it proceeds from one spin state and is forbidden from the other, so the unreactive state is kinetically protected and the correlated spin state persists on timescales relevant to biochemical processes. SCRP-dependent chemistry has been proposed as a mechanism underlying avian magnetoreception and photosynthetic electron transfer^10,11,12,13,14,15^. In bird cryptochromes, flavin adenine dinucleotide (FAD) is photoreduced by tryptophan to form an SCRP across ∼20 Å^11,16^. In the radical pair mechanism (RPM), direct back electron transfer to the FAD ground state proceeds only from the singlet state, so if singlettriplet interconversion occurs within the coherence time, flavin fluorescence becomes field-dependent. The sign of this effect reports the pair’s birth multiplicity: a magnetic field increases fluorescence for a singlet-born pair and decreases it for a triplet-born one^2,3,17^. Such magnetic field effects (MFEs) have been observed in cryptochromes, where the singlet-born FAD-tryptophan pair brightens with field^4,11^, in photolyases^11,18,19^, and in free flavin solutions with reductants^17,20,21^. Whether cryptochrome’s magnetosensitive biological signaling derives from this RPM remains unsettled^4^.

Light-Oxygen-Voltage (LOV) domains are ubiquitous flavoproteins in plants, fungi, bacteria, and the human gut microbiome^22,23,24^. The LOV2 domain of *Avena sativa* phototropin 1 (AsLOV2) is a blue-light photoreceptor widely used in optogenetics^6,25,26,27^. In AsLOV2, blue light drives covalent bond formation between FMN’s C4a and the sulfur atom of cysteine 450 (C450). This pho-toadduct is widely thought to form through a radical pair rather than an ionic intermediate, on the basis of time-resolved EPR and ultrafast spectroscopy, though the pair itself has not been directly observed^28,29^; computational work^30^ and deuterium kinetic isotope effects on bond formation^31^ suggest that it forms by hydrogen-atom transfer from the cysteine to the flavin. The photoadduct is the protein signaling state, with subsequent J*α* helix undocking as a consequence of cysteine-flavin bond formation^32,33,34,35^. This places AsLOV2’s radical pair directly on its signal transduction pathway.

Wild-type (wt) AsLOV2 shows no MFE at fields up to 25 mT^5^ and has been classified as magnetically inactive. In every flavoprotein in which an MFE had been demonstrated prior to this study, the radical partners lie far apart (≥15 Å) and are thus weakly coupled, and none carry a reactive cysteine. The FMN–cysteine photoadduct motif, in which the radicals sit ∼4.4 Å apart and are strongly coupled, is the conserved, canonical arrangement across the LOV family^6^. These two facts are not independent. Weak coupling, which arises when the radicals are far apart, leaves a radical pair sensitive at low field; the ∼4.4 Å distance in AsLOV2 that allows covalent bond formation also raises the dipolar coupling to hundreds of MHz, pushing the minimal field for an MFE out of the range previously surveyed. Across the flavoproteins studied so far, low-field sensitivity and cysteine-binding signaling competence are therefore anti-correlated for a structural reason. The classification of AsLOV2 as magnetically inactive is thus an artifact of the field range examined: as we show here, strong coupling and Δ*g*-driven interconversion push the MFE onset to high field, and hyperfine-driven mixing between distant, weakly coupled radicals is not the only route to flavoprotein magnetosensitivity.

MagLOV (and, similarly, iLOV)^3^ is constructed to avoid the flavin-cysteine photoadduct, replacing C450 with a proline and using a distant (∼15 Å) native tryptophan as the electron donor^36^, producing a cryptochrome-like tryptophan-flavin radical pair driven by hyperfine asymmetry^2,5^. Using site-directed spin-labeling, we measure that MagLOV lacks the same conformational change as wt AsLOV2 (S.6, Fig. S7). MagLOV carries five substitutions relative to wt AsLOV2 (C450P, L496V, Q513K, G528K, D540M). Besides removing the reactive cysteine, Q513K removes the conserved glutamine that gates the wild-type photocycle^32^, so its failure to undergo the wild-type conformational response has two independent structural grounds. MagLOV displays a large fluorescent MFE, up to about 30% at 20 mT^2,5^, and it and similar flavoproteins have been demonstrated as in-cell reporters^2,5,9^. Because MagLOV does not form the cova-lent FMN–cysteine photoadduct, its MFE is not directly connected to a known downstream signaling event in a way that AsLOV2 is.

MagLOV and wild-type AsLOV2 are therefore opposite cases: MagLOV has a large MFE but does not form the known signaling photoadduct, while AsLOV2 forms the signaling photoadduct^28,29,33,34,35,37^ but was reported magnetically in-active^2,5,36^. Coherent radical-pair spin dynamics and biological signal transduc-tion had never been observed in the same protein. Because AsLOV2’s photocycle is well-understood^38,39^, its MFE can be tied directly to downstream biochemical events: photoadduct formation drives the J*α* helix undocking^32,33,34^ which ac-tivates the natural kinase or alternate optogenetic effector domains, so an MFE offers a magnetic handle on function.

We report an MFE on wt AsLOV2 that emerges only at high field, acting directly on chemical bond formation and, thereby, on protein signaling state. Fluorescence experiments show that the wt AsLOV2 MFE grows with field, plateauing near 1 T, while the MagLOV MFE is already large at 20 mT. UV–Vis experiments with and without a magnetic field reveal an MFE on FMN–cysteine bond formation itself. We model the field dependence to show how strong electron coupling and *g* -factor asymmetry together produce the observed MFE. These results establish a quantum signal transduction mechanism in a wild-type protein and a design constraint for engineering magnetosensitive ones. Given the ubiquity of LOV domains, the coupling of high-field magnetosensitivity to signaling may be widespread.

## 2 Results

### 2.1 Fluorescence MFE in wild-type AsLOV2 emerges only above 75 mT

We began by measuring the magnetic field dependence of fluorescence under continuous blue-light excitation in both MagLOV and AsLOV2 at fields ranging from 20 to 1300 mT using an electromagnet for field control (setup shown in Fig. S2, processing details in S.1 and Fig. S1). For wild-type AsLOV2, an MFE emerged at ∼75 mT and grew in magnitude with magnetic field strength: 1% at 150 mT (Fig. 3(d)), 5% at 650 mT, and 7% at 1300 mT (Fig. 1(c); full field dependence in Fig. 3(d)). AsLOV2 therefore exhibits a fluorescence MFE, but only at field strengths well above those previously surveyed for flavoprotein magnetosensitivity, where earlier work had found no MFE^2,5^.

**Figure 1:**
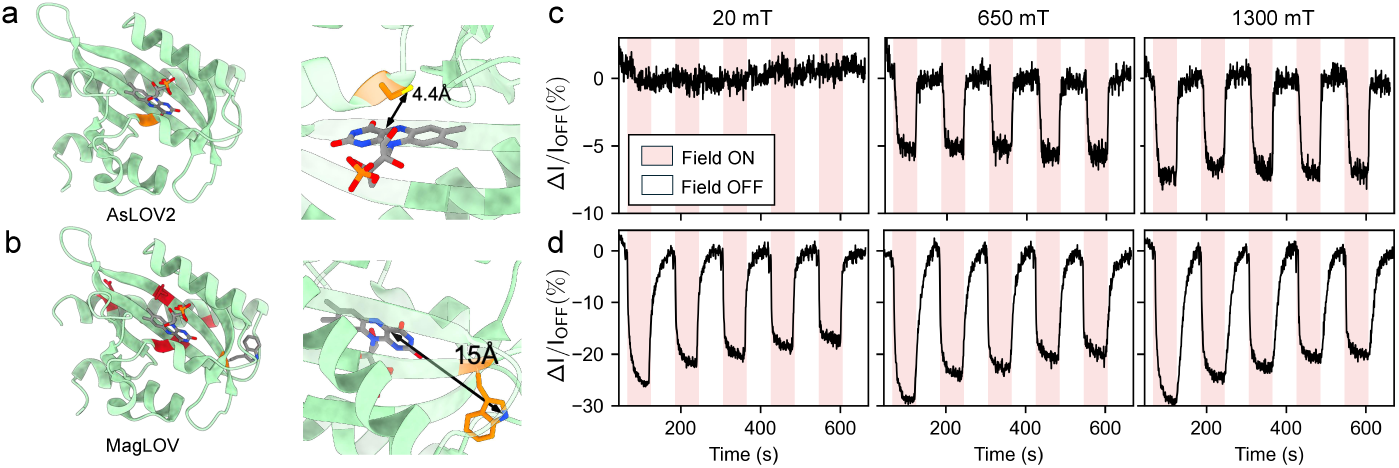
(a) A protein structure of AsLOV2 (PDB 2v1a) with the covalently binding cysteine and flavin highlighted in a blow-up subsection. **(b)** A protein structure of MagLOV (PDB 2v1a) with the 5 residues mutated from wild-type AsLOV2 highlighted in red. The tryptophan key for the electron transfer step and the flavin are highlighted in a blow-up subsection. **(c)** AsLOV2’s MFE grows in magnitude with increasing field, with the field-on fluorescence unchanged at 20 mT and decreasing by 5% at 650 mT and 7% at 1300 mT. In the red shaded regions, a magnetic field is turned on. In response to this field, a change in normalized fluorescence is observed. For a full description of AsLOV2’s MFE field-dependence see Fig. 3. **(d)** In contrast, MagLOV exhibits an MFE with a much lower magnetic field onset: fluorescence intensity decreases by a relatively constant extent across fields, 25% at 20 mT, 30% at 650 mT, and 30% at 1300 mT. Decreases in the MFE size over an experiment have been observed previously^2^ and are due to damage of the MagLOV sample by the laser.

**Figure 2:**
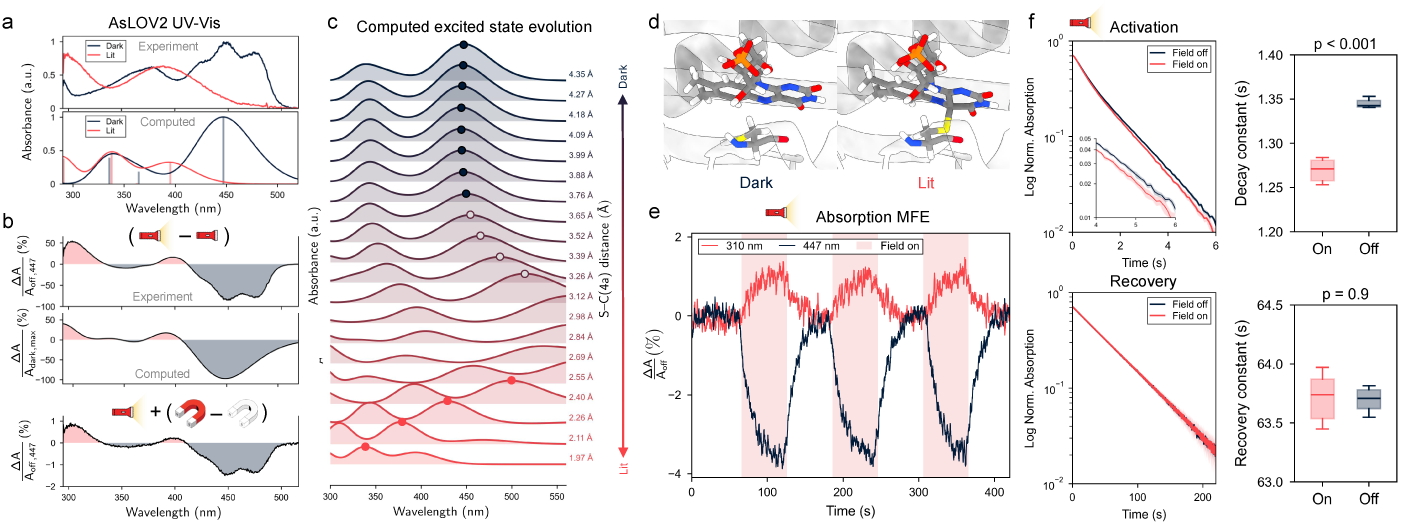
Magnetic field effect on AsLOV2 bond formation. **(a)** UV–Vis absorp-tion spectra for dark and lit AsLOV2 states, linked to cysteine bond formation and conformational response, as shown by AsLOV2 photoactivation. Computed spectra (TDDFT, *ω*B97X-D4/def2-TZVP; Methods M.11) are shifted by *−*0.375 eV, which places the dark-state band at 447 nm; bars mark individual transitions. **(b)** Normalized difference in UV–Vis spectrum for light vs. dark conditions (top) and field on vs. off conditions (bottom) with light on, showing that the field-induced changes mirror the light-induced changes. the computed lit-dark difference is shown for comparison. **(c)** Computed evolution of UV–Vis absorption spectra with the S–C(4a) distance go-ing from the unbound to the bound state along an interpolated path between the dark and lit structures (circles: lowest bright band). **(d)** Structures of AsLOV2’s C450 and FMN chromophore in the dark, unbound state and the lit, bound state. **(e)** Time-dependent magnetic field effects (650 mT field) on the equilibrium absorption at 310 nm (peak for lit state) and 447 nm (peak for dark state), showing opposite sign MFEs that confirm the field-induced shift towards the lit state. **(f)** Activation (top) and recovery (bottom) of cysteine bond as measured by absorption at 447 nm for field on (650 mT) and off conditions. Activation experiments were recorded with the light turned on at *t* = 0, and recovery experiments were recorded with the light turned off at *t* = 0 following 5 minutes of photoexcitation. Activation from the dark to lit state is sensitive to whether the field is applied, with a significantly different decay constant (p *<* 0.001), while recovery back to the dark state is not sensitive to a magnetic field (p = 0.9).

**Figure 3:**
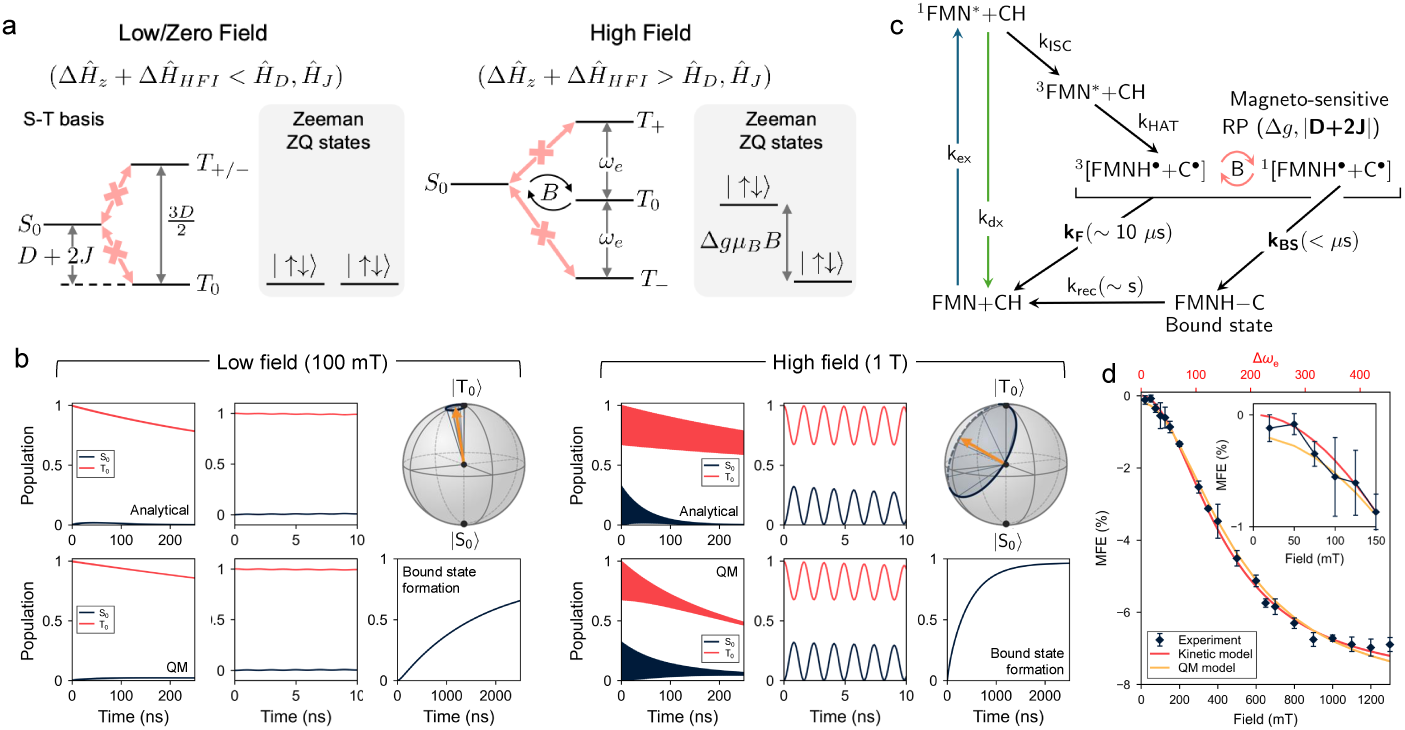
Mechanism behind AsLOV2 magnetic field effects. **(a)** Low and high field regimes (dipolar basis vs Zeeman basis) allowing interconversion only at fields where the asymmetry in the Zeeman interaction is significant compared to the dipolar interaction. **(b)** Interconversion of *T*0 and *S*0 at low (left) and high (right) field using both the analytical model in Eq. 3 and quantum mechanical calculations in Spinach under Haberkorn kinetics. Top right shows the Bloch sphere representation of the singlet–triplet interconversion, with the effective Hamiltonian axis the orange arrow and the state evolution in blue. Bottom right shows the bound state population as a function of time modeled in Spinach, showing significantly faster bound state formation at high field. **(c)** AsLOV2 photocycle kinetics to model the AsLOV2 fluorescence MFE. The bolded parameters (*|D* + 2*J |, k_f_ , k_BS_*) were varied to fit the experiment data. **(d)** Simulation of fluorescence MFE field dependence using kinetic model described in text and quantum mechanical simulations. Full rates and model for both kinetic and quantum mechanical calculations described in S.7 and S.8 and Table 1.

This high-onset field dependence originates in the short FMN-cysteinyl radi-cal separation (∼ 4.4 Å), where dipolar and exchange couplings far exceed those of the FMN/FAD–tryptophan radical pairs typical of known magnetosensitive flavoproteins. The same proximity that sets the field threshold also lets the radical pair react directly, forming the FMN–cysteine bond of the photoadduct. This is a chemical reactivity native to wt AsLOV2 and absent from previously studied magnetosensitive flavoproteins (mechanism in Sec. 3.3)^4,11,17,18,40^. This high-onset field dependence of the fluorescence intensity is readily recapitulated on the benchtop using inexpensive switchable neodymium permanent magnets that switch from ∼40 mT to above 300 mT (S.3, Fig. S3), making high-onset measurements broadly accessible.

MagLOV shows a strong MFE that, unlike AsLOV2’s, is readily saturated at low field. Previous studies reported a fluorescence MFE in MagLOV of up to 30% at 20 mT with limited field dependence over 10 − 50 mT (descendant MagLOV2 is similar)^41^, and our measurements at 20 mT reproduce this with a ∼25% MFE (Fig. 1(d)). Across 650 and 1300 mT the MagLOV MFE stays nearly constant, changing by only 5% between 20 and 1300 mT (Fig. 1(d)).

This near-independence is the signature of a hyperfine-driven system: once the Zeeman splitting (560 MHz at 20 mT) exceeds the hyperfine difference, *S*_0_ − *T*_±_ interconversion is suppressed while the *S*_0_ − *T*_0_ gap is field-independent, so that further increasing the field has little effect. MagLOV reaches this regime because its reactive cysteine is mutated to proline, so the electron donor is a distant (∼1.5 nm) tryptophan^36^ rather than the nearby cysteine of AsLOV2 (Fig. 1(a,b)). In both AsLOV2 and MagLOV, the effect of the magnetic field decreases fluo-rescence as expected for a triplet-born radical pair, though through different mechanisms between the two variants (Sec 3.3), and opposite to the increase in fluorescence intensity seen with the singlet-born radical pair in cryptochromes^4^.

### 2.2 Magnetic field modulates formation of the covalent photoadduct

Previous work identified the FMN–cysteine radical pair formation as a hypoth-esized, though silent, intermediate in the formation of the covalent C-S bond that produces the AsLOV2 signaling state^28,29,37^. If the fluorescence MFE we observed arises from this radical pair, then the chemical bond formation step itself should be magnetic field dependent. A fluorescence MFE, while diagnostic of a radical pair, reports only on excited-state branching, that is, on how spin state partitions the radical pair between decay channels; it does not establish that the field changes the amount of covalent product formed.

We hence turn to UV-visible absorption (UV–Vis) spectroscopy above the field threshold established by the fluorescence results. The dark and lit states of AsLOV2 have well-resolved UV–Vis signatures (Fig. 2(a))^34^ from disruption of FMN’s aromaticity due to C-S covalent bond formation (Fig. 2(c)): the dark state has maximal absorption at 447 and 473 nm and the lit state at 380 nm and *<* 300 nm (Fig. 2(a)), giving a direct readout of the signaling-state population. Time-dependent density functional theory (TDDFT) calculations on an FMN–C450 model reproduced both spectra with a single uniform shift (Fig. 2(a,b); Methods M.11, S.10). The 447 nm band is a flavin-localized *π* → *π*^∗^ transition (93% of the hole density on the isoalloxazine) that requires conjugation through C4a. It disappears once C4a becomes sp^3^ in the adduct, whose near-UV absorption (computed at 291 and 338 nm) accounts for the lit-state gain at 310 nm (Fig. S13). Along a computed path from the dark structure to the adduct (Fig. 2(c)), the 447 nm band is essentially the same at the 4.39 and 3.75 Å S–C4a distances of the two dark-state C450 rotamers (position within 1 nm, intensity within 20%) from the crystal structure (PDB 2V1A)), while the lit-state bands appear only at bonded distances (≤2.1 Å). The 447 and 310 nm signals therefore report the populations of the unbound dark state and covalent adduct, respectively, not a redistribution between pre-reactive rotamers.

To measure absorption intensity changes experimentally, we measure UV-Vis absorption MFE using a setup built into the electromagnet pole gap (Fig. S4; Methods) while cycling the field between 650 mT and off, mirroring the flu-orescence MFE protocol. Application of the 650 mT field produced a UV–Vis lineshape shift consistent with increased lit-state population, demonstrated by the similarity of the lineshape changes for photo-induced bond formation and field-modulated bond formation (Fig. 2(b)). To resolve this population shift quantitatively, we monitored two wavelengths with maximum contrast between the dark and lit states – 447 nm (dark, unbound state) and 310 nm (lit, bound state) – during repeated cycling on and off of the magnetic field. The 447 nm absorbance decreased under the applied field, indicating a decrease in the pop-ulation of the dark, unbound state, while absorption at 310 nm correspondingly increased (Fig. 2(e)). The 650 mT magnetic field, therefore, shifts the equilib-rium population from the unbound towards the bound flavin states. By tracking the photoadduct UV–Vis signature, we measure the field effect on product for-mation directly, namely the population of the covalent FMN–cysteine bond that defines the signaling state, as opposed to the fluorescence field effect that reports on the population of the fluorescent (ground) FMN state. Transient UV–Vis ab-sorption is an established method in the flavoprotein MFE literature, but it has been used to follow the radical pair rather than the photoadduct; the photolyase effect was read at 460 and 510 nm over 1–9 *µ*s at 39 mT^18^, as were the cryp-tochrome^4,11^ and free-flavin^20^ measurements, while the engineered-protein and in-cell studies read emission^2,3,5,9^. To our knowledge, this is the first demon-stration of an MFE measured directly on functional covalent bond formation from a spin-correlated radical pair.

Magnetic field effects on FMN–cysteine bond formation also appear in the activation kinetics from dark to lit state, but not in the reverse recovery pro-cess. We monitored the absorbance at 447 nm following the onset of blue-light excitation with the field off and with a 650 mT field on, and fit the data to an exponential decay to extract the activation rate, *k_act_* (Methods M.8). Acti-vation in the 650 mT field was faster than without the external field showing 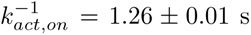 and 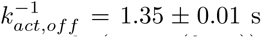 with a significant *p*-value of *p <* 0.001 across 6 different trials (Fig. 2(f,top)), consistent with the magnetic field increasing the rate of bond formation by rendering the radical pair spin dynamics more favorable towards bond formation. This rate enhancement corresponds to a 6.7 ± 0.6% kinetic MFE (Methods M.9), in approximate agreement with the fluorescence MFE observed at the same field of 5% (Fig. 1(c)). As a control against systematic artifacts, we measured the recovery kinetics (FMN–cysteine thermal bond cleavage following light shut-off), which does not proceed through a spin-correlated radical pair mechanism. The recovery time constant 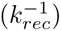, extracted by exponential fit, showed no significant difference between field-on and off (*p* = 0.9; Fig. 2(f,bottom)). An effect on both the forward and the reverse steps would indicate a change in the sample conditions rather than in the spin chemistry, since thermal reversion is a ground-state, water-gated process with no radical pair intermediate^32^. At 350 mT, both equilibrium population shifts and activation showed the same field response as in 650 mT, but with a smaller effect, as expected at lower field (Fig. S6). The magnetic field therefore acts specifically on the bond formation pathway through the spin-correlated radical pair mechanism.

### 2.3 *g*-factor asymmetry (Δ*g*) overcomes strong dipolar coupling at high field

Singlet-triplet interconversion in an SCRP is driven by asymmetry between the two unpaired electrons’ environments; in the limit of identical environments, only relaxation (no coherent interconversion) occurs. For MagLOV and the avian cryptochrome, this asymmetry arises from differential hyperfine interac-tions, and the ∼1.5 − 2 nm inter-radical distance makes the dipolar coupling weak (∼few MHz) and negligible to first order^2,36^. At low magnetic field, the hyperfine difference efficiently mixes *S*_0_ with all three triplet states, and as the field is increased, the Zeeman energy separates the *T*_±_ states from *S*_0_ and sup-presses the hyperfine-driven mixing, producing the conventional low-field MFE.

For wt AsLOV2 the geometry differs in kind. The conserved reactive cysteine is located in the close vicinity of the isoalloxazine ring, so the radical pair is formed in van der Waals contact. In the room-temperature (293 K) dark-state structure (PDB 2V1A), C450’s sulfur is refined as two alternate rotamers, 4.39 Å from FMN C4a at 0.70 occupancy and 3.75 Å at 0.30, with light shifting the population towards a shorter distance attributed to the covalently bound state^35^. The point-dipole coupling

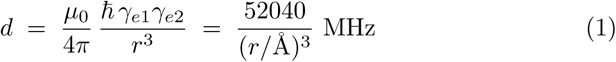

is therefore 616 MHz in the major rotamer and 990 MHz in the minor one, and 728 MHz is the weighted average. Either value exceeds the hyperfine couplings of both radicals by more than an order of magnitude^42,43^, so the dipolar term sets the zero-field level structure.

This strong dipolar coupling exceeding 600 MHz suppresses triplet-singlet interconversion in the low-field limit (Fig. 3(a,b)), since the low-field eigenstates of the dipolar Hamiltonian are *S*_0_*, T*_0_*, T*_+_*, T*_−_ (or *T_x_*, *T_y_* and *T_z_*, depending on the symmetry of the system). The FMN-cysteinyl radical pair is distinct from a FMN-tryptophan radical pair in that it has a much greater Δ*g* than for organic radicals: *g*_iso,FMN_ = 2.0032^44^, while cysteinyl has a wide range of simulated *g*-factors (accounting for hydration) ranging from 2.03 to 2.10^45^ and a measured isotropic of *g*_iso,cyst_ = 2.028^46^. All are *>* 1% different from *g*_iso,FMN_. Therefore, as the magnetic field increases, the electron Larmor frequency asymmetry Δ*ω_e_* = *ω_e_*_1_ − *ω_e_*_2_ grows, with Δ*ω_e_* = 34.7 MHz at 100 mT and Δ*ω_e_* = 347 MHz at 1 T, enabling mixing between *S*_0_ and *T*_0_. In AsLOV2 the *T*_±_ states remain significantly separated in energy at both low field (by dipolar coupling) and high field (by Zeeman energy), so their coupling to *S*_0_ is inefficient regardless of initialization and occupancy, as evidenced by the lack of low field onset MFE (See S.7 for more details). This mechanism resembles the Δ*g*-effect, which leads to high-field (generally but not always *>* 1 T)^21,47,48^ changes in recombination and back-reaction yield in photoresponsive proteins^21,49^, quantum dots^47,50^ and organic radical reactions^48,51,52^, but these systems generally lack the strong inter-radical spin coupling present in AsLOV2^21^.

We next model the observed fluorescence MFE on the basis of the Δ*g*-effect with spin physics calculations of a radical pair. We restrict our analysis to the two-level basis spanned by the *S*_0_−*T*_0_ manifold. Within this manifold, we use the spin Hamiltonian consisting of the Zeeman Hamiltonian, 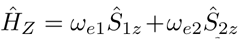, the secular dipolar Hamiltonian, 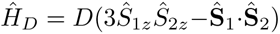 with 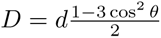, and the exchange Hamiltonian *H̭_J_*= −2*J***S̭**_1_· **S̭**_2_, yielding

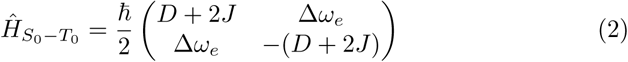

where 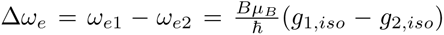, with *µ_B_* the Bohr magneton, scales linearly with field strength. D here is the secular, orientation-dependent, part of the dipolar coupling. From the form of this Hamiltonian, it is clear that the off-diagonal Δ*ω_e_* mixes *S*_0_ and *T*_0_, with the degree of mixing set by the ratio of the off-diagonal elements Δ*ω_e_* to the diagonal elements *D* + 2*J*. Here *θ* is the angle between the S*γ*–C4a dipolar axis and the field. *D*(*θ*) runs from −*d* at *θ* = 0 through zero at 54.7^◦^ to +*d/*2 at *θ* = 90^◦^, the most probable orientation of the protein at any given time. For a *T*_0_-initialized radical pair state, the probability of generating a singlet state radical pair evolves with time *t* as

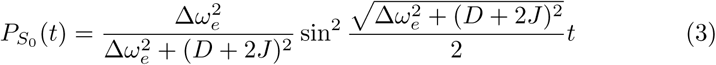

with the maximum singlet probability 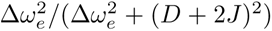 reached when 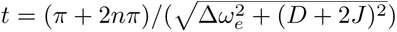 The singlet–triplet interconversion is shown in the low (100 mT) and high (1 T) field limits in Fig. 3(b) for this analytical equation. We also performed full quantum mechanical calculations of the spin dynamics in the Haberkorn kinetics model in the fast motion limit using the Spinach program package^53^ (see S.8 for details), allowing us to explicitly include relaxation and the influence of the *T*_±_ states. Both sets of results, shown in the left panels, show that the maximum singlet population and interconversion rate are field dependent, with larger singlet populations observed as the field increases. The frequency of the singlet-triplet oscillatory interconversion, seen in the zoom-in, is identical between the analytical and quantum mechanical models, confirming that the analytical approach appropriately captures these oscillations at short times.

This interconversion is also shown on the Bloch sphere spanning the |*T*_0_⟩ − |*S*_0_⟩ states, where the orange arrow indicates the effective Hamiltonian axis, and the state vector procession around the effective Hamiltonian is shown in blue. Significantly, the field-induced difference in singlet populations calculated from the integrated singlet yield in the Haberkorn model manifests as a different rate and maximum of bound state formation (Fig. 3(b), bottom), supporting the experimentally observed changes in Fig. 2 where higher magnetic field increases the rate and equilibrium population of the bound state.

Using this understanding of singlet-triplet interconversion for strongly cou-pled radicals, we constructed a quantum-mechanically informed kinetic model of the wt AsLOV2 MFE (Fig. 3(c)). The model couples the AsLOV2 photo-cycle and the coherent singlet-triplet interconversion defined by Eq. 3, along with decoherence time, *T*_2_, to capture the finite lifetime of the spin correlation.

This state structure is inspired by similar models for work on wt and mutant AsLOV2^2,28,54^. We use known values for the g-factors of FMN^44^ and cysteinyl radicals^46^, and for the radiative decay rate, *k_dx_*, and the intersystem crossing rate, *k_ISC_*^29^. *k_ex_* is estimated based on the laser power used, while the rate of recovery from the photoadduct to the fluorescent ground state, *k_rec_*, is mea-sured via UV–Vis (Fig. 2(d)). The rate of SCRP formation via hydrogen-atom transfer (*k_HAT_* ) is set to be 5 ∗ 10^5^ s^−1^ based on previous estimates^29,31^.

This leaves three free parameters: the *S*_0_ −*T*_0_ energy gap, |*D* +2*J*|, the spin-nonselective escape rate to the ground state, *k_f_* , and the rate of FMN–cysteine bond formation, *k_BS_*, all directly related to the radical pair dynamics. The singlet-triplet interconversion is described by a pair of rates *k_S_*_−→_*_T_*and *k_T_* _→−_*_S_* chosen to reproduce the equilibrium singlet pop<u>ulation</u> 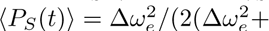 Since the interconversion rate 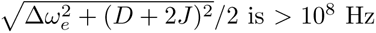 from the effect of (*D* + 2*J*) alone, i.e. even at zero field, the coherent precession is faster than the kinetic decay rates from the radical pair. Therefore, relevant dynamics can be captured using time-averaged populations, without resolving individual oscillations. For more details on the kinetic modeling, see S.7.

The fit of this kinetic model to the experimental field dependence of the fluorescence MFE is shown in Fig. 3(d), with best fit parameters |*D* +2*J*| = 500 MHz, *k_f_* = 8.8 ∗ 10^4^ s^−1^, and *k_BS_* = 20.5 *µ*s^−1^. This value of *k_BS_* is consistent with the literature, which proposes that bond formation from the singlet state should proceed at a timescale faster than *µ*s^29^. Similarly, the *S*_0_ − *T*_0_ energy gap of |*D* + 2*J*| = 500 MHz is consistent with known AsLOV2 FMN–cysteine distances of 3.7 – 4.4 Å^35^ - given that the population-weighted dipolar coupling (in the point dipole approximation) between the two structures is 728 MHz, and most probable value of *D* is *D*_⊥_ = *d/*2 = 364 MHz. This sets the value of *J* to be of the order of +70 MHz or −850 MHz, depending on the sign of |*D* + 2*J*|.

Neither the sign nor the magnitude of J can be determined from the fitting. Furthermore, these fit values for the *S*_0_ − *T*_0_ energy gap |*D* + 2*J*| is likely a lower bound arising from accounting only for the secular Dipolar Hamiltonian term. Future quantum chemical calculations could extract, or at least constrain, the sign and value for the exchange coupling in order to put better bounds on *J*.

This spin dynamic modeling does not yield entirely unique set of parameters; there is some correlation between *k_f_* , *k_BS_*, and |*D* + 2*J*|. We scanned the parameter space by fixing each parameter to the value used in the fit above while varying the other two to determine the goodness of fit. These results, shown in Fig. S9, show that there is a strong correlation between |*D* + 2*J*| and *k_BS_*, while *k_f_* is approximately fixed at just under 10^5^ s^−1^. This correlation has a physical meaning - larger |*D* + 2*J*| results in lower singlet population, so producing equivalent bound state yield requires a faster rate of bound state formation from the singlet state. While this means that we cannot uniquely fix |*D* + 2*J*| and *k_BS_*, the geometric constraints on the dipolar coupling^35^ and the approximate timescale of *k_BS_*^29^ do place reasonable restrictions on likely value ranges for each parameter; |*D* + 2*J*| on the scale of several hundred MHz and *k_BS_* on the order of 10 − 100 *µ*s^−1^.

We also varied each parameter in the kinetic model independently, and found that the magnetic field dependence is strongly influenced by the parameters that directly impact the radical pair dynamics - |*D* + 2*J*|, Δ*g*, *k_f_* , *k_BS_*, and *T*_2_ - while the other kinetic rates in the photocycle have minimal impact on the resulting field dependence (Fig. S10). In particular, the decoherence time of *T*_2_ is correlated with the spin nonselective escape rate *k_f_* , as their product sets the escape fraction, i.e. radical pairs that do not undergo bond formation. Δ*g*, on the other hand, is correlated with |*D* + 2*J*| and *k_BS_*, as these three values determine the maximum singlet population and probability of bond formation from the singlet state at a given field.

To confirm these results, we simulated the singlet yield (a proxy for covalent bond formation and MFE) via a full quantum mechanical simulation in the fast motion regime (Section S.8). The Haberkorn kinetic model^55^ was used, where the kinetic rate of the bound state formation, *k_BS_*, was set to the rate of formation of the singlet state of the radical pair, and the escape rate, *k_f_* , was set to the kinetic rate of the triplet state deactivation, while the magnetic field effect was assumed to be proportional to the relative change in singlet yield as a function of field compared to the zero-field value. The results (Fig. 3(d)) are in excellent agreement with the analytical model. We find that only when *T*_1*e*_ is small enough, i.e. comparable to the radical pair lifetime set by *T*_2_, such that significant *T*_±_ populations build up during the radical pair lifetime do the singlet-triplet interconversion and MFE vary compared to the analytical model (Fig. S11(b)). These quantum-mechanical simulations also show that the shape of the field dependence for the singlet yield is not impacted by the initialization of the triplet state population (Fig. S11(c)), justifying the approach used in the kinetic model that takes into account only the *S*_0_ − *T*_0_ manifold.

We note that this model uses a single value of |*D* + 2*J*|. In reality, *D* is modulated by the protein rotation and *J* is not independently known, since spin dynamics lie in the slow-motion regime. This regime is characterized by ratios of times and interaction magnitudes: the rotation correlation time of AsLOV2 (molecular weight 17 kDa) in aqueous solution is of the order of *τ_c_* ∼ 5 − 10 ns, the dipolar coupling modulation amplitude is 3*d/*2 ∼ 1000 MHz, and the radical pair lifetime/bond formation is of the order of 50 − 1000 ns. A full quantitative analysis of the magnetic field effect in AsLOV2 aimed at extracting exact values for |*D* + 2*J*|, *k_f_* , and *k_BS_* would thus require treatment with the stochastic Liouville equation, together with independent determination of the *J* value, which will be the focus of future studies.

## 3 Discussion and Conclusion

The FMN–cysteine radical pair in AsLOV2 differs from previously studied photo-excited SCRPs in protein systems. The combination of the radicals’ close spacing and strong coupling with the distinct *g* -factors of cysteine and FMN pro-duces a high-field (*>* 75 mT) MFE with strong field dependence over the range of 20 − 1300 mT, reaching 7% at maximum. The same close proximity that creates strong dipolar coupling is normally, but not always^56,57^, taken to suppress magnetosensitivity. Close proximity, on the other hand, also enables FMN–cysteine bond formation from the radical pair, providing a direct route from a quantum spin state to a biochemical signaling state. In this sense, the radical pair functions as a two-state molecular qubit consisting of the *S*_0_ singlet and *T*_0_ triplet states that coherently interconvert and whose interconversion can be controlled. In fact, this radical pair dynamics goes beyond presenting a qubit system to achieving quantum transduction by coupling the qubit state directly with distinct biochemical outcomes (bond formation vs. back reaction to the ground state).

The field-dependent singlet-triplet interconversion within a radical pair is the most plausible mechanism of MFE in this system, and the only mecha-nism proposed to describe the MFEs in flavin systems. The existence of a wt AsLOV2 MFE provides strong evidence that a previously inferred^28,29^ but never observed spin-correlated radical pair lies on the AsLOV2 photoadduct pathway. The strong dipolar and exchange coupling, together with the Δ*g*-driven mech-anism, have made it inaccessible to transient EPR^28^, which requires resolvable, weakly coupled spin populations. Direct spectroscopic observation was there-fore largely precluded by the very coupling regime that defines the system. The SCRP has also proven silent in transient IR spectroscopy^29,37^. The MFE pro-vides the diagnostic that transient EPR and IR spectroscopy have not: a field effect on both fluorescence and product formation, detected by the absorption MFE, establishes the SCRP without requiring its direct detection. The fields required to observe AsLOV2-type MFEs and probe the initialization of a SCRP are accessible with inexpensive switchable permanent magnets (Fig. S3). Not readily observable ODMR with wt AsLOV2 (S.9) is consistent with this picture: using *D* = 728 MHz and *J* of either +70 or −850 MHz, the X-band linewidth is *>* 1 GHz, which is not easily addressable under physiological conditions and with established instrumentation.

Beyond establishing the radical pair, the AsLOV2 MFE has implications for similar behavior in other protein systems. Its high-field onset means that protein SCRPs operating in regimes where Δ*g*-driven interconversion dominates over hyperfine-driven interconversion were overlooked at the lower magnetic fields (*<* 30 mT) that dominate the protein MFE literature. Given that LOV domains are ubiquitous across plants, fungi, bacteria and the human gut microbiome^22,23,24^, other naturally occurring magnetosensitive proteins with high-field MFEs may have eluded detection. We hypothesize that a wide range of wt LOV domains have different spin and kinetic parameters and therefore different shapes and magnitudes of the magnetic field response. For proteins that do proceed via radical pairs, fitting an MFE vs field curve is simultaneously a measurement of its coupling and a screen for magnetic addressability. Overall, these results suggest that high-field MFE measurements of both fluorescence and UV-Vis absorption can be used as a tool for probing photoadduct formation pathways in other proteins and testing whether they proceed via radical pairs.

The specific geometry of AsLOV2 is not necessarily optimal for a large MFE at any given field. The field onset and field dependence are determined by the inter-radical distance and the *g*-factors of the participating radicals, both of which are tunable through site-directed mutagenesis or directed evolution. Our absorption MFE provides a probe with potential for high-throughput analysis that mechanistically links together magnetosensitivity and covalent reactivity, which could be used to evolve altered MFEs while requiring FMN–cysteine re-activity. Mutants with weaker dipolar coupling or larger Δ*g* could shift the field onset below 50 mT, where readily generated laboratory fields could en-able magnetically tuned bond formation in optogenetic or magnetogenetic sys-tems^6,25,26,27,58,59^ or biological sensing via multiplexed MFEs^2^. As with the directed evolution of MagLOV, mutations affect not only the radical pair partners but also the kinetics of the photocycle and may control MFE characteristics^5,36^. The design constraint this work exposes is that the two properties must be engineered against each other: moving the radicals apart lowers the onset field but removes the reactivity, which is the trade that produced MagLOV^5^ and the protein-qubit sensors built from it^2,60^. What AsLOV2 shows is that the other side of that tradeoff, with nearby radicals that are directly reactive, does not imply magnetic silence, only a different magnetic regime. This engineering space is the design space for biological quantum devices: protein systems in which a single qubit, embedded in a controlled molecular environment, directs a single biochemical decision.

The coherent interconversion of AsLOV2’s SCRP is biologically relevant at Earth’s field, since *T*_0_–*S*_0_ interconversion is suppressed and bond formation in a triplet-initialized spin state cannot occur prior to decoherence. This creates a natural “quantum barrier” to bond formation, an effect analogous to Pauli spin blockades in double quantum dots^61^. The rate of bond formation is therefore limited by SCRP decoherence, which is sensitive to protein environment and rotational correlation time and therefore could be tunable to some degree by nature. The lifting of this quantum barrier can be readily achieved at high magnetic field, which is an effect that should be considered when photoactivating LOV domains in high fields (such as in NMR or EPR)^33,62^.

In summary, AsLOV2’s magnetosensitive SCRP links an FMN–cysteine bond-forming mechanism directly to a well-characterized wild-type protein signaling state. The emergence and field dependence of a previously unseen high-field MFE in AsLOV2 are captured by the interplay of strong dipolar coupling and an unusually low-onset Δ*g*-effect. The same magnetic field modulates both the protein fluorescence and the UV–Vis absorption spectrum that reports on FMN–cysteine bond formation preceding conformational activation. This mechanism links coherent spin dynamics to the formation of the FMN–cysteine photoadduct and LOV signaling state. The FMN–cysteine MFE has potential to add mag-netic control to optogenetic systems or optical sensing platforms that already in-tegrate LOV domains. The result establishes a magnetosensitive FMN–cysteine bond-forming reaction, and indicates that magnetically reactive protein systems operating in the strongly coupled, Δ*g*-driven regime may be more widespread than the prevailing focus on hyperfine-driven MFEs has suggested.

## Methods

Protein expression, purification, and spin-labeling protocols were derived from Maity et al. 2025^34^.

### **M.1** Protein expression and purification

DNA fragments encoding the LOV2 domain of *Avena sativa* phototropin 1 (residues 403–546), the MagLOV variant (C450P, D540M, Q513K, L496V, G528K)^5^, and K413C mutants of each were purchased from GENE UNIVER-SAL or GenScript, cloned into pET-26b(+) via NdeI (5^′^) and XhoI (3^′^) restric-tion sites with a C-terminal 6×His tag. Plasmids were transformed into *E. coli* BL21 (DE3) and plated on LB agar containing kanamycin (0.05 mg/mL). A single colony was used to inoculate 10 mL LB + kanamycin, grown at 37 ^◦^C with shaking (220 rpm, 18 h). The primary culture was used to inoculate 1 L TB autoinduction medium containing kanamycin and incubated for ∼16 h at 18 ^◦^C in the dark with shaking.

Cells were harvested by centrifugation at 5,000 *g* for 15 min at 4 °C, resuspended in lysis buffer (20 mM Tris-HCl, 500 mM NaCl, pH 8.0, 5 mM *β*-mercaptoethanol, 1 mg/mL lysozyme), incubated 30 min at 4 °C, lysed by sonication (60 × 10 s pulses with 10 s intervals), and clarified at 11,000 rpm for 30 min at 4 °C. Lysate was supplemented with 6 mg FMN in 1 mL water (15 min, room temperature, 15 rpm, dark), then combined with 10 mL Ni-NTA resin (2 h, 4 ^◦^C, 15 rpm, dark). Resin was washed three times with 40 mL wash buffer (20 mM Tris-HCl, 500 mM NaCl, 40 mM imidazole, 5 mM *β*-mercaptoethanol, pH 8.0). Protein was eluted with 10 mL elution buffer (as wash buffer, with 500 mM imidazole) and further purified by size-exclusion chro-matography (HiLoad^TM^ 16/600 Superdex^TM^ 75 pg, GE Healthcare; NGC^TM^ system, Bio-Rad; SEC buffer: 20 mM Tris-HCl, 150 mM NaCl, pH 8.0). Con-centrations were determined by absorbance at 447 nm (AsLOV2) or 443 nm (MagLOV) using *ε* = 13,800 M^−1^ cm^−1^.

### **M.2** Site-directed spin-labeling

Purified AsLOV2 K413C was reduced overnight at 4 ^◦^C with 10 mM DTT, then desalted (PD-10, GE Healthcare). MagLOV K413C does not tolerate overnight DTT exposure and was used without reduction. Both samples were reacted with a 10-fold molar excess of MTSL (1-Oxyl-2,2,5,5-tetramethyl-Δ^3^-pyrroline-3-methyl methanesulfonothioate) overnight at 4 ^◦^C with gentle shaking in the dark. Excess label was removed by size-exclusion chromatography (HiLoad^TM^ 16/600 Superdex^TM^ 75 pg, equilibrated with 20 mM Tris-HCl, 150 mM NaCl, pH 8.0). Final spin-labeled samples were concentrated to ∼400 *µ*M.

### **M.3** Inter-radical distances

Inter-radical distances were extracted using ChimeraX-1.10.1 from the AsLOV2 crystal structure (PDB 2V1A) 293 K dark state^35^. The AsLOV2 radical pair distance was taken as the FMN C4a to C450 SG distance. C450 SG is refined in two alternate positions, 4.389 Å from C4a at 0.70 occupancy and 3.746 Å at 0.30, which gives point-dipole couplings *d* = 616 and 990 MHz and an occupancy-weighted *d* = 728 MHz (effective separation 4.15 Å). The MagLOV radical pair distance was approximated by the FMN C4a to Trp491 CE2 distance on the same structure, since MagLOV’s substitutions do not affect the relevant back-bone geometry. The estimated MagLOV distance is consistent with simulated values from Antill and coworkers^36^.

### **M.4** X-band CW EPR

Continuous wave (CW) EPR spectra were acquired at 293 K on a Bruker EMX X-band spectrometer with a dielectric cavity featuring an optical window (ER 4123D). Samples (10 *µ*L) were loaded into 0.66 mm i.d. quartz capillaries (Vit-roCom). Photoexcitation used a nominally 450 nm blue laser (peak ∼ 460 nm as measured using OceanOptics USB 2000+; nominal power 70 mW, lower at the sample due to optical-window diffraction grating; Laser Components USA). Spectra were recorded in the dark and under continuous 460 nm illumination at a center field of 0.35 T, microwave frequency 9.8 GHz, microwave power 2 mW, modulation amplitude 0.5 G, modulation frequency 100 kHz, sweep width 150 G, averaged over 5 scans.

### **M.5** X-band fluorescence MFE

Fluorescence measurements were performed at 293 K using a Bruker ElexSys E580 X-band EPR spectrometer and Bruker ER073 electromagnet. Samples (60 *µ*L) in 3 mm quartz EPR tubes (Wilmad) were photoexcited by a 460 nm blue laser (70 mW, Laser Components USA), focused through a Thorlabs F220SMA-532 collimator into a BFH37-600 multimode fiber (600 *µ*m diameter) positioned ∼1 cm from the sample. A breadboard between the Helmholtz coils held a 2-inch, 65 mm focal-length plano-convex lens, a Thorlabs 495 nm longpass filter, and a 50 *µ*m Thorlabs fiber connected to a USB 2000+ OceanOptics spectrophotometer. Each data point reflects the fluorescence spectrum acquired in 600 ms; the 480 − 600 nm region was integrated to quantify fluorescence. The field was cycled on/off in 1-minute periods for 2-min cycles, with 5 cycles per experiment (11 min total), beginning and ending field-off (e.g. 1). AI was used to assist the generation of the Python script used to extract integrals from the fluorescence data.

The AsLOV2 MFE vs. field curve (Fig. 3d) was acquired by switching the field intensity during each off cycle, in 2 minute cycles as described above, in a randomly ordered sequence of 20 field strengths (41 min per run, triplicate). The fields used are 20 mT, 50 mT, 75 mT, 100 mT, 125 mT, 150 mT, 200 mT, 300 mT, 350 mT, 400 mT, 500 mT, 600 mT, 650 mT, 700 mT, 800 mT, 900 mT, 1000 mT, 1100 mT, 1200 mT, and 1300 mT. Sample stability over this duration was verified by repeating the 1300 mT measurement for 20 cycles with consistent response.

### **M.6** Spin dynamics simulation and MFE fitting

The field dependence of the AsLOV2 fluorescence MFE was modeled using a kinetic-model fit with the spin dynamics treated within the *S*_0_–*T*_0_ manifold and time-averaged.

We used the kinetic scheme shown in Fig. 3(c), which couples the AsLOV2 photocycle to singlet-triplet interconversion within the FMN–cysteine spin-correlated radical pair. The radical pair was initialized in the *T*_0_ state. Triplet-singlet inter-conversion was modeled as a pair of rates *k_S_*_→_*_T_* and *k_T_* _→_*_S_* chosen to reproduce the time-averaged singlet population

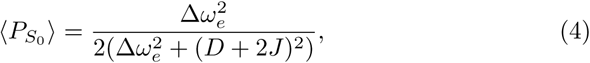

where Δ*ω_e_* is the field-dependent Larmor frequency difference between the two radicals and *D* + 2*J* is the *S*_0_–*T*_0_ gap with *D* the dipolar coupling and *J* the exchange coupling. An additional decoherence state *RP_dec_* captured the loss of spin correlation through *T*_2_ relaxation. Simulations were run in MATLAB using *fmincon* constrained nonlinear least-squares minimization routine, with *k_ex_*, *k_dx_*, *k_ISC_*, *k_HAT_* , *k_rec_*, *R*_2_, and the *g*-factors of FMN and cysteine fixed, and |*D* + 2*J*|, *k_f_* , and *k_BS_* as free parameters. The full rate table and additional fitting details are given in S.7.

### **M.7** Tabletop fluorescence MFE

Tabletop fluorescence measurements were acquired at room temperature using a Magnetpro Multipurpose Switch Magnet (660 lbs lifting strength) clamped to an optical table (Fig. S3). Magnet stability during switching was verified by recording mirror-reflected light intensity over 5 on/off cycles. Field at the magnet surface was ∼300 mT (on) / 40 mT (off), measured with a LATNEX MF-30K gaussmeter. The reflective magnet surface was painted black. Samples (50 *µL*) in rectangular borosilicate capillaries (VitroCom, 0.4 ×4 ×50 mm) were photoexcited at 45^◦^ with the same 460 nm laser used in M.4, with intensity adjusted to provide fluorescence without heating. Fluorescence collection optics (plano-convex lens, 495 nm longpass filter, 50 *µ*m Thorlabs fiber, USB 2000+ OceanOptics) were identical to M.5; field cycling and quantification were also as in M.5.

### **M.8** UV–Vis absorption MFE

UV–Vis absorption measurements used a deuterium lamp from a Hamamatsu High Power UV–Vis Fiber Light Source source coupled to a Thorlabs FOFMS-UV In-line fiber filter mount (250-450 nm). The FOFMS-UV was positioned between the poles of the Bruker EMX X-band electromagnet using a 3D-printed holder. A 100 *µ*M AsLOV2 sample in a 1 cm plastic cuvette was illuminated from above by a compact blue laser (70 mW, 460 nm, Laser Components USA) through a BFH37-600 600 *µ*m multimode fiber held in place by a 3D-printed cap. An estimated 3 mW was transmitted to the sample. Transmitted white light was collected by a 50 *µ*m Thorlabs fiber connected to a USB 2000+ OceanOptics spectrophotometer. Background was acquired with a water-filled cuvette. AI was used to assist the generation of the Python script used to extract time-dependent wavelength information from the absorption data.

#### Equilibrium experiments

Field cycling matched the fluorescence MFE pro-tocol (1-min on/off, 2-min cycles, 5 cycles, 11 min total). Each data point captured 90 ms of the full 290–550 nm spectrum at 0.4 nm resolution.

#### Activation and recovery kinetics

Activation was measured by recording 480 nm absorbance following onset of blue-light excitation; recovery was measured following light shutoff after 60 s of irradiation. Each data point captured 90 ms of absorbance. Field-on and field-off experiments used one undisturbed sample, with experiment order randomized. Activation time courses were fit to a single exponential decay,

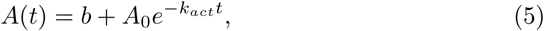

where *A*(*t*) is the absorbance at time *t*, *A*_0_ is the absorbance change between dark and lit states, *b* is the baseline absorbance at *t* → ∞, and *k_act_* is the activation rate. Recovery time courses were fit to

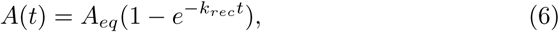

where *A_eq_* is the equilibrium dark-state absorbance and *k_rec_* is the recovery rate. Each condition was measured across 6 independent trials. Statistical significance was assessed by paired *t*-test (5 degrees of freedom). The 650 mT recovery rate difference Δ*k_rec_* was −0.15 ± 0.32%.

### **M.9** MFE quantification

Fluorescence MFE was defined as Δ*I/I_off_* = (*I* − *I_off_* )*/I_off_* , where *I* is the fluorescence intensity and *I_off_* is the field-off intensity from a background spline fit to the field-off periods (Fig. S1). UV–Vis MFE (Fig. 2(b,e)) at wavelength *λ* was defined analogously as Δ*A/A_off_* . For Fig. 2(f), the percent change in UV–Vis activation and recovery time constants (Δ*τ* ) was calculated as

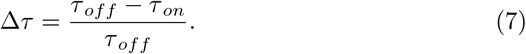

Error bars on the on and off rates were 95% confidence intervals (*t*-test, 5 degrees of freedom), and error in Δ*τ* was propagated as

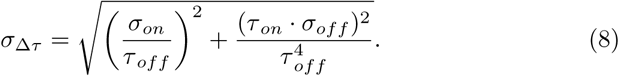

Spline-fitting details are in SI S.1.

### **M.10** Optically detected magnetic resonance

ODMR was acquired on a Bruker EMX X-band spectrometer with a Bridge 12 microwave power source (up to 10 W). MagLOV samples (100 *µ*M in 3 mm cap-illaries) were placed in a tuned cylindrical dielectric resonator. The fluorescence collection optics matched M.5 (plano-convex lens, 495 nm longpass, 50 *µ*m fiber, USB 2000+). The field was swept 3254.4–3454.4 G in 66 s with microwaves at 9.407 GHz, 36 dBm (4 W). Fluorescence acquisition was synchronized with microwave onset and sweep start. MagLOV was not temperature-controlled.

AsLOV2 ODMR (500 *µ*M) was attempted at 4 ^◦^C using a Variable Tempera-ture Unit, sweeping 2920–3720 G in 300 s at 9.80 GHz with powers between 4 and 10 W; no measurable contrast was observed (see S.9).

### **M.11** Electronic-structure calculations

*Model and path.* The cluster model (63 atoms, charge −2) contains the com-plete FMN cofactor (isoalloxazine, ribityl chain and dianionic 5^′^-phosphate) and the C450 side chain, with the backbone capped as an amine and an aldehyde. The dark endpoint (C450 thiol, oxidized FMN; S–C4a 4.35 Å) and the lit endpoint (C4a–S adduct, N5-protonated FMN; S–C4a 1.97 Å) were taken from constrained-optimized dark (2V0U) and lit state (2V0W) AsLOV2 PDB struc-tures^35^. Intermediate structures were generated by screw interpolation. Along this path the S–C4a distance shortens to form the final bonded state, and the thiol proton moves from S to N5.

*Absorption spectra.* All calculations used ORCA 6.0 with RIJCOSX (def2/J) and CPCM (*ε* = 4) for the protein environment. Singlet excitation energies and oscillator strengths (20 roots) were computed with TDDFT in the Tamm–Dancoff approximation (TDA) from a restricted *ω*B97X-D4/def2-TZVP refer-ence. Stick spectra were broadened with Gaussians in energy (*σ* = 0.17 eV). A single uniform shift of −0.375 eV, which places the dark-state S_1_ band (3.149 eV, 394 nm) at the experimental 447 nm, was applied to every image; the lit state was not adjusted separately. Transitions were assigned from their leading excitation amplitudes and natural transition orbitals (NTOs). Orbital charac-ter was quantified by integrating |*ϕ*|^2^ over isoalloxazine, cysteine, ribityl and phosphate fragments (nearest-atom partition).

## Acknowledgments

The authors acknowledge support through the NSF CHE CMI Grant 2505621, and NIH Grant Number R01AG05605. S. Han, S. Maity, and S. Chaudhuri ac-knowledge support by the NIH MIRA **R35GM136411** grant and the Deutsche Forschungsgemeinschaft (DFG, German Research Foundation) as a part of Ger-many’s Excellence Strategy EXC2033 390677874 RESOLV. W. Salvia acknowl-edges the National Institute for General Medical Science (NIGMS), through Northwestern University’s Chemistry of Life Processes (CLP) Predoctoral Train-ing Program **T32GM149439**, as well as the National Institutes of Health Training Grant (**T32GM140995**) through Northwestern University’s Molecular Bio-physics Training Program. J. Straub acknowledges the NRSA from the National Institute of General Medical Sciences under award number 1F32GM161114-01A1. The content is solely the responsibility of the authors and does not necessarily represent the official views of the National Institutes of Health.

The authors would like to thank Quentin Stern, Hoang Le, Martyna Judd, Maria Ingaramo and Rebecca Hayward for valuable discussions, Amani Al-ghamdi and Franz Geiger for helping with optical setups and providing com-ponents, Jinlei Cui for instrumental support and Savannah Vennard, Megan Vettoretti, Jonah Rand, and Ojas Sharma for help with protein purification.

## Supplementary Information

See associated file **SupplementaryInformation.pdf**

