## Supplementary Information for "Quantum transduction from electron spin state to signaling in wild-type AsLOV2"

### Supplementary Information: Quantum transduction from electron spin state to a signaling state in a wild-type LOV photoreceptor

William Salvia*[1], Joshua Straub*[1], Shiny Maity[1], Alexey Bogdanov[2], Subhajyoti Chaudhuri[1], Ruixian Han[1], and Songi Han[1]

[1]Department of Chemistry, Northwestern University, Evanston, IL 60208, USA
[2]Department of Chemical and Biological Physics, Weizmann Institute of Science, Rehovot 7610001, Israel

November 18, 2026

*These authors contributed equally to this work.

### Supplementary Information

#### S.1 Spline fit for MFEs in fluorescence and UV–Vis data

Using fluorescence spectra from the OceanOptics spectrometer, we integrated the region between 480 and 600 nm at each timepoint. We then plotted these integral values vs. time. To control for baseline drift and initial saturation period in AsLOV2 and MagLOV magnetic field effects, we implemented a background spline fit using manually selected points at the final field off point before the field was turned on for each cycle, following Abrahams et al.[1]. The background spline fit was then subtracted from the experimental data, and the processed data was plotted as described in section M.9. The steps in this processing are shown below for an AsLOV2 fluorescence MFE dataset taken at 650 mT.

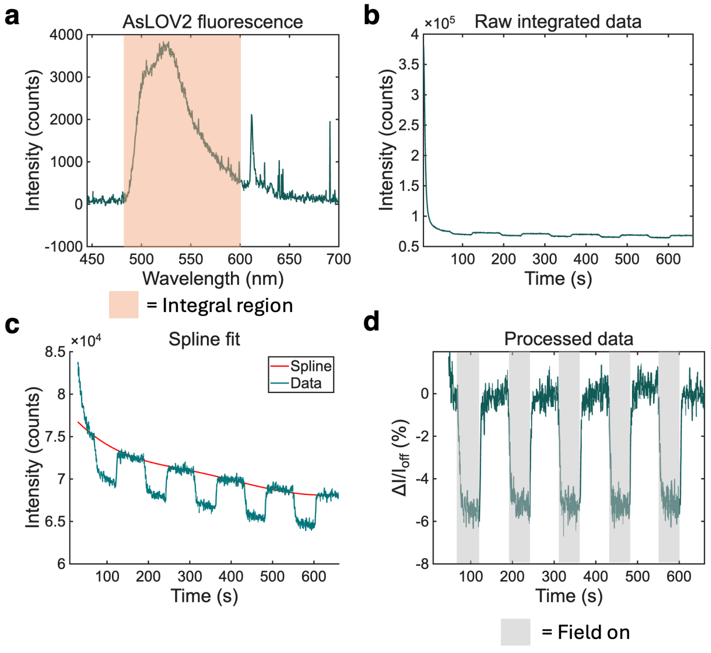


Figure S1: Processing of MFE data. **(a)** AsLOV2 fluorescence spectrum under blue light illumination, with integration region shaded in red. **(b)** Raw integrated fluorescence counts. **(c)** Spline fit to the background (field off) fluorescence. **(d)** Processed MFE data, plotted as $\Delta I/I_{off}$, with field on regions marked in grey.

#### S.2 X-band optical setup for fluorescence MFE

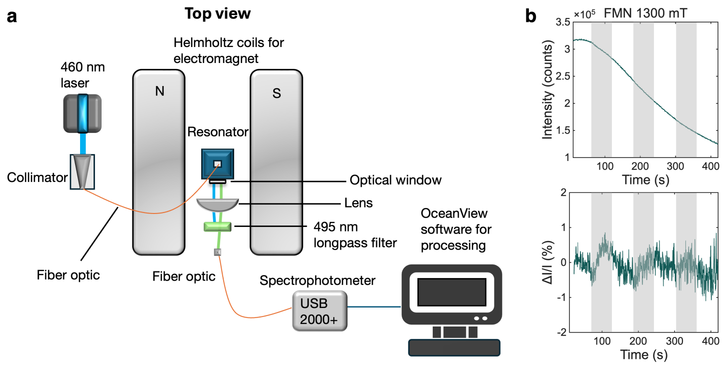


Figure S2: Fluorescence MFE setup. (a) Schematic used for fluorescence MFE measurements.. (b) FMN control on setup. Top is the raw data for FMN fluorescence, and bottom is processed data after spline fitting. Shaded gray regions indicate field on (1300 mT).

To confirm that the observed fluorescence MFE in MagLOV and AsLOV2 for our setup was from the magnetically altered fluorescence of the protein, we conducted a control experiment on FMN alone. Without an electron donor in the sample, FMN should not show any magnetic field effects on its fluorescence. We thus ran a MFE experiment on a sample of 400 $\mu$M FMN at 1300 mT for three field on-off cycles following the same protocol as used for the AsLOV2 and MagLOV samples. The data is shown in Fig. S2(b), and shows no MFE for FMN. Note that the fluctuations in the intensity are not periodic with magnetic field, and are likely caused by sample diffusion with respect to the beam path.

#### S.3 Tabletop optical setup for fluorescence MFE

The AsLOV2 MFE is accessible for experimental study without the need for high-field electromagnets. On a tabletop setup, we used a switchable magnet (see M.7 for details), which can be purchased for \$60 on Amazon, to observe the AsLOV2 MFE. The magnitude of the MFE on this setup was ∼3% at 300 mT with the magnet switched "on" and 40 mT with the magnet switched "off" (Fig. S3(c)), a comparable size to the 300 mT field on the X-band fluorescence set-up using an electromagnet.

To control for any systematic error from magnet field switching, we secured a 1.5 inch thick wooden block to the front of the magnet to reduce the field strength (with the field on) experienced by the same AsLOV2 sample to 40 mT as measured by the gaussmeter. If the change in fluorescence was due to factors such as changed sample position, we would expect a change in the fluorescence even at this low field value. No change was observed (Fig. S3(b)).

An additional control was an 400 $\mu$M FMN sample in buffer (20 mM TRIS, 150 mM NaCl, pH 8.0), expected to have no MFE. When placed flush against the magnet, no MFE was observed at 300 mT (Fig. S3(d)).

These experiments establish that AsLOV2's MFE is reproducible on different setups. This is a low-budget setup ($<$ \$1000 total) for others interested in studying AsLOV2-like MFEs using fluorescence. This setup also has the methodological advantage of selecting only for high field-onset MFEs, since 40 mT $->$ 300 mT is not a significant change for hyperfine-driven MFEs like MagLOV's.

This setup also provides a probe of a magnetosensitive LOV domain's SCRP initialization, if a LOV domain progresses through a FMN-cysteine covalent bond. AsLOV2 has suppressed singlet-triplet interconversion at low fields and increased interconversion at high fields (the reverse of MagLOV, where high field suppresses singlet to $T_{\pm}$ interconversion), but both proteins show the same negative MFE. This is because MagLOV's radical pair singlet state undergoes relatively fast back electron transfer to the ground state, such that higher singlet populations lead to faster recovery to the ground state, and thus more fluorescence. In contrast, the AsLOV2 singlet state forms an FMN-cysteine bond, a long-lived dark state with slow relaxation to the ground state, such that higher singlet population leads to less fluorescence. The same MFE sign therefore reports on different mechanisms underlying the MFE. In the case of a positive MFE in a LOV domain, that would suggest that the SCRP initialization is singlet-borne, and does not progress through an ISC prior to SCRP formation.

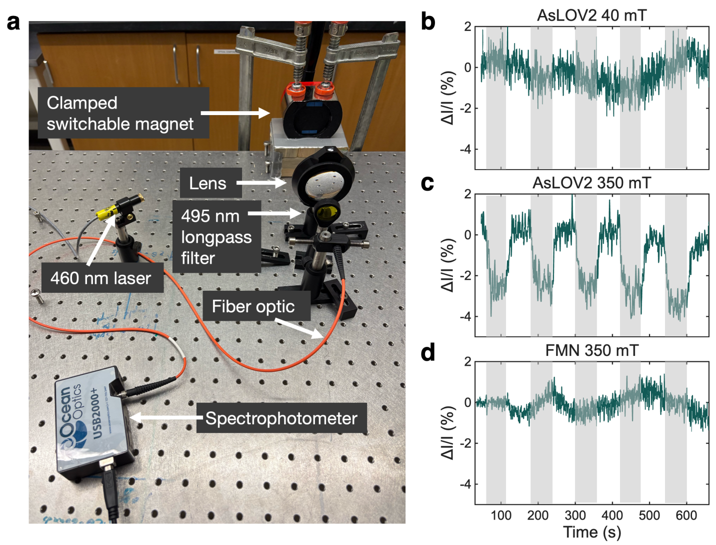


Figure S3: Tabletop MFE setup. **(a)** Picture of setup used to record table fluorescence MFE. **(b)** AsLOV2 fluorescence MFE when the a wooden block is placed between the magnet and the sample, so that the field on period has a magnetic field of ∼40 mT and the field off period has roughly zero field. **(c)** AsLOV2 fluorescence MFE when the sample is directly on the magnet, generating a field of ∼300 mT for the magnet on period and ∼40 mT for the magnet off period. The magnitude of the MFE agrees with the value measured at 350 mT on the setup using the electromagnet. **(d)** FMN fluorescence MFE with a 400 $\mu$M FMN sample placed directly on the magnet. For (b)-(d), grey regions indicated periods of field on.

#### S.4 UV–Vis absorption MFE setup

For the absorption MFE experiment at an equilibrium between cysteine bound and unbound AsLOV2, we illuminated the sample using the setup shown in Fig. S4 such that the photoactivation rate matched the recovery rate and the UV–Vis spectrum remained at a steady-state, with visible cysteine-unbound absorption peaks at 447 and 473 nm (Fig. S5).

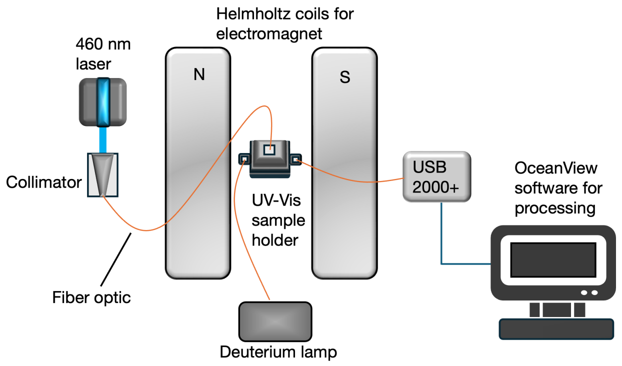


Figure S4: Setup for UV-Visible absorption MFE measurements as viewed from above.

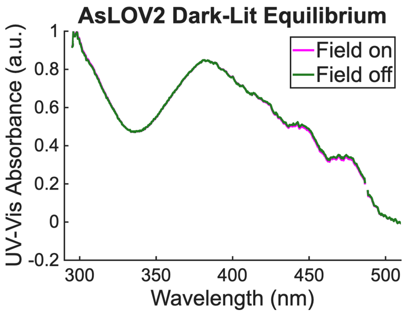


Figure S5: UV–Vis absorption spectrum under equilibrium conditions for both field on (650 mT) and field off conditions. A comparison to the spectrum shown in Fig. 2(a) shows that features of both the dark and lit state are present. The difference between these spectra is shown in Fig. 2(b).

#### S.5 UV–Vis absorption MFE on AsLOV2 activation kinetics at 350 mT

We used the same protocol as the 650 mT experiment, except that we positioned the UV–Vis sample holder at a spot in the X-band magnet such that the field at the sample was weaker (350 mT). UV–Vis absorption was then measured at 447 nm as a function of time after the laser was initially turned on (activation), or after the laser was turned off following 60 s of irradiation (recovery) over 7 experiments with the same untouched sample. The average time constant of activation, fit to a monoexponential (see main text for details), is significantly different with field on/off: $2.65 \pm 0.05$ s with the field on and $2.72 \pm 0.04$ s with the field off ($p = 0.03$) (Fig. S6(a)). This is a $2.6 \pm 2.3$ % MFE for the time constants, using the mean and error equations from Methods M.8.

This value is consistent with the fluorescent MFE at 350 mT (Fig. 3(d)). The differences in activation time constant between 350 mT and 650 mT datasets are due to different setup conditions, since the activation rate of AsLOV2 is highly dependent on the intensity of laser light for photoactivation, and the laser and collimator was positioned differently both days.

The difference between field on and field off recovery time constants was $1.0 \pm 1.8$ %, non-significant ($p = 0.22$) (Fig. S6(b)). Differences between recovery constants in these conditions compared to Fig. 2(d) are likely due to temperature effects from different laser induced heating across different setups or ambient temperature differences in the room. A significant activation magnetic field effect and non-significant recovery effect is consistent with the 650 mT data (Fig. 2(f)).

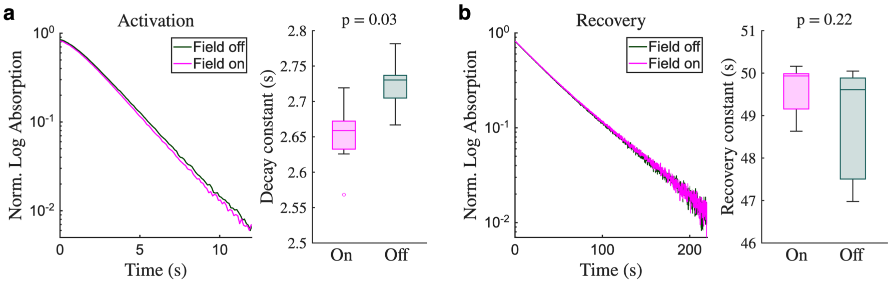


Figure S6: AsLOV2 kinetic MFE at 350 mT. **(a)** Average activation for field on and field off (left) and decay constants (right) across 7 independent trials, showing a statistically different decay constant ($p = 0.03$). **(b)** Average recovery for field on and field off (left) and decay constants (right) across 7 independent trials, showing no statistical difference in recovery constants ($p = 0.22$).

#### S.6 Conformational response of AsLOV2 and MagLOV

By adding an MTSL spin label to AsLOV2 with site K413 mutated to a cysteine, we found that it exhibits a very large change in X-band CW EPR lineshape compared to previous work (compare to less-responsive spin-labeled positions in Maity et al. 2023[2] and Maity et al. 2025[3]). This large change in lineshape is indicative of increased conformational restriction of the spin label upon protein conformational change by a Microscopic Order Macroscopic Disorder (MOMD) model[4]. We consider this change in lineshape to be a proxy for the unique AsLOV2 unfolding mechanism that can be monitored at room temperature with a standard X-band EPR spectrometer. When MagLOV is labeled at the same site, it has a very similar spectrum in the dark state. However, we do not see a change in lineshape upon photoactivation like that of wild-type AsLOV2's, just a decrease in intensity. The observed change is likely due to a baseline feature that likely corresponds to MagLOV's photobleaching and subsequent aggregation. These data indicate that MagLOV does not respond to light with an AsLOV2-like conformational response. This is not a trivial result despite MagLOV's lack of the flavin-binding cysteine, since LOV domain VIVID is known to enter a signaling state even when the cysteine is removed[5].

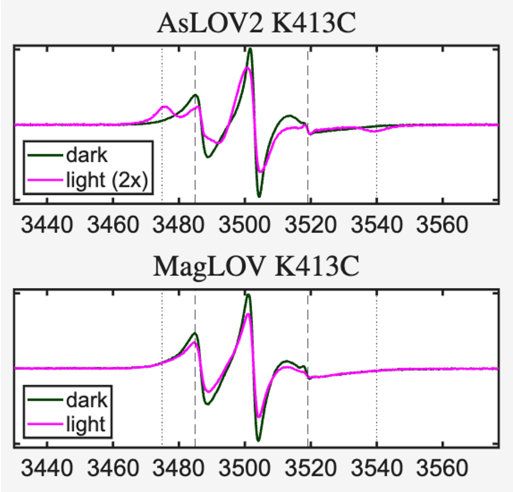


Figure S7: Comparison of K413C mutated and MTSL labeled AsLOV2 and MagLOV CW EPR spectrum in the dark and exposed to a 460 nm laser. The light state spectrum for AsLOV2 K413C is scaled 2-fold so that the lineshape is easily compared to the dark state.

#### S.7 Kinetic modeling of AsLOV2 fluorescence MFE

The kinetic model used to fit the AsLOV2 fluorescence MFE is shown in Fig. S8. Blue-light excitation of the FMN ground state generates a singlet excited state, which undergoes intersystem crossing to a triplet excited state. Hydrogen atom transfer from C450 to FMN forms the spin-correlated radical pair (RP) in a triplet state, with magnetic field–dependent singlet-triplet interconversion governed by Eq. 3. The decoherence state $\mathrm{RP}_{\mathrm{dec}}$ captures the loss of spin correlation through $T_2$ relaxation. RP (and $\mathrm{RP}_{\mathrm{dec}}$) decay to the ground state in a spin-nonselective manner or proceed through the spin-selective covalent FMN–cysteine bond formation from the singlet state (yielding the long-lived signaling state, which slowly recovers to the ground state).

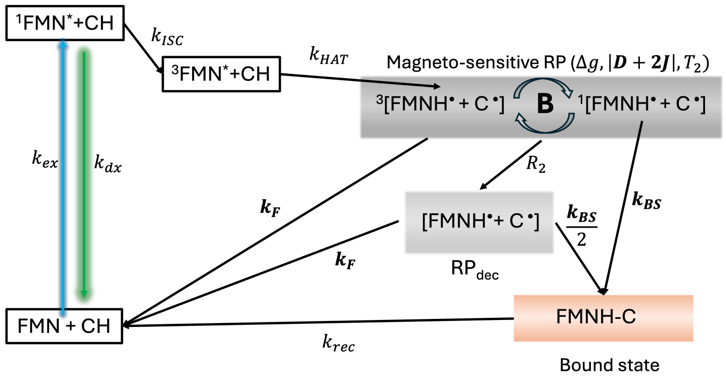


Figure S8: Photocycle used for kinetic modeling of fluorescence MFE. In comparison to the photocycle shown in Fig. 3(c), we add in the additional state $\mathrm{RP}_{\mathrm{dec}}$, to account for $T_2$ relaxation of the radical pair.

We initialize the radical pair in the $T_0$ state. Following intersystem crossing, the triplet is spin-polarized with unknown populations of $T_-$, $T_0$, and $T_+$, but the $T_\pm$ states never efficiently interconvert with $S_0$: at low fields interconversion is blocked by the strong dipolar coupling, and at high fields by the large Zeeman energy difference. Since $T_{1e}$ is expected to exceed $T_2$ for the radical pair, we assume negligible mixing of the triplet states during the SCRP lifetime. Different initial triplet polarizations are absorbed into the rate of SCRP formation in the model. Note that there could be zero-field mixing of $S_0$ and $T_\pm$ when $|D/2-2J|$ is small, allowing the differential hyperfine interaction to efficiently mix these states. However, if there was significant $S_0 - T_\pm$ mixing at zero field, we would see a fluorescent MFE as we increased the field on the order of tens of mT, where the Zeeman splitting would then decouple any $S_0 - T_\pm$ that was allowed. The lack of any MFE at these low field values means that it is appropriate to neglect this potential zero-field mixing.

We can estimate the rate of excitation $k_{ex} = I_0 \epsilon_A l \log 10$, where $I_0$ is the incident laser intensity in Einstein $\mathrm{L}^{-1}\ \mathrm{s}^{-1}$, $\epsilon_A \sim 10^4\ \mathrm{M}^{-1}\ \mathrm{cm}^{-1}$ for FMN at 450 nm[6], and $l \sim 0.5$ cm is the estimated depth of light propagation in the sample at the region of the optical window. With a measured laser power output from the fiber optics of $\sim 1$ mW, we get an excitation rate of $k_{ex} \sim 4.25$ s$^{-1}$.

As described in the main text, we fit the experimental MFE using this model with $k_{ex}$, $k_{dx}$, $k_{ISC}$, $k_{HAT}$, $k_{rec}$, $R_2$, and the $g$-factors for FMN and cysteine fixed, and $|D+2J|$, $k_f$, and $k_{BS}$ as free parameters. $k_{ex}$ was estimated as described above, while $k_{dx}$, $k_{ISC}$[7], and the $g$-factors for FMN[8] and cysteine[9] were taken from literature. $k_{rec}$ was measured with UV–Vis. $k_{HAT}$ was estimated to be $5 * 10^5$ s$^{-1}$ based on previous transient absorption measurements[7,10]. $R_2$ was estimated at 1 MHz. With $R_2$ fixed, $k_f$ is determined only relative to it: the fit returns the escape fraction $k_f/R_2 = 0.088$ ($R_2$ from $10^5$ to $10^7$ s$^{-1}$ gives $k_f$ from $8.8 \times 10^3$ to $8.8 \times 10^5$ s$^{-1}$ with identical fit quality. Constraints were placed on the ground state population based on the degree of fluorescence saturation observed. The rates for singlet-triplet interconversion were set to establish the equilibrium singlet population $\langle P_S(t) \rangle = \frac{\Delta\omega_e^2}{2(\Delta\omega_e^2+(D+2J)^2)}$ within 10 ns, given that the timescale for singlet-triplet interconversion is at minimum $(D+2J)/2 > 100$ MHz, as described in the main text. This equilibrium assumption is appropriate since this interconversion is still an order of magnitude greater than the fastest kinetic rate out of the radical pair state ($k_{BS}$).

The simulation shown in Fig. 3(d), resulting from the best fit to the experimental data, used the parameters listed in Table 1.

Table 1: Rate constants and spin parameters for the AsLOV2 fluorescence MFE simulation. Rates and $|D+2J|$, $R_2$ in Hz; $g$-factors dimensionless.

| Parameter | Value |
|---|---|
| $k_{ex}$ | 4.25 |
| $k_{dx}$ | $6 \times 10^8$ |
| $k_{ISC}$ | $6 \times 10^8$ |
| $k_{HAT}$ | $5 \times 10^5$ |
| $k_f$ | $8.8 \times 10^4$ |
| $k_{BS}$ | $2.05 \times 10^7$ |
| $k_{rec}$ | 0.02 |
| $g_{FMN,iso}$ | 2.0032 |
| $g_{cyst,iso}$ | 2.028 |
| $R_2$ | $1 \times 10^6$ |
| $\|D+2J\|$ | $500 \times 10^6$ |

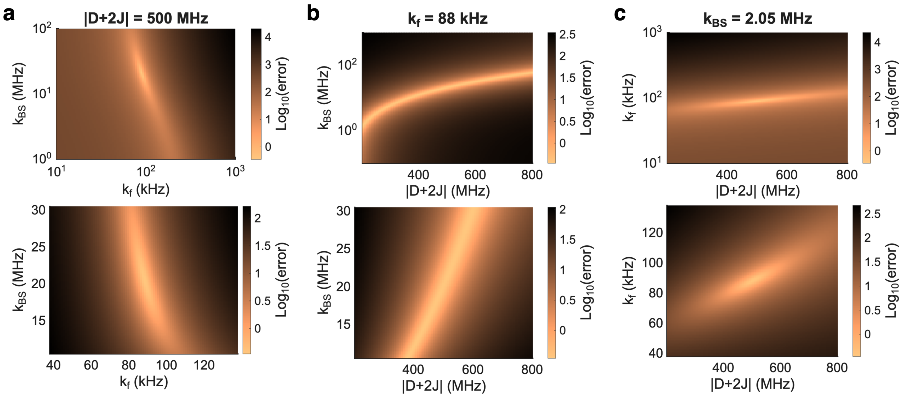


Figure S9: Kinetic MFE simulations showing how varying the three fit parameters ($|D+2J|, k_f, k_{BS}$) impacts the resulting fit. For each plot, one of the three fit parameters was fixed at its best fit value - (a) fixes $|D+2J|$, (b) fixes $k_f$, and (c) fixes $k_{BS}$. The top row varies $k_f$ and $k_{BS}$ over several orders of magnitude and is log-scale for the kinetic parameters, while the bottom plot is a zoom in on a linear scale.

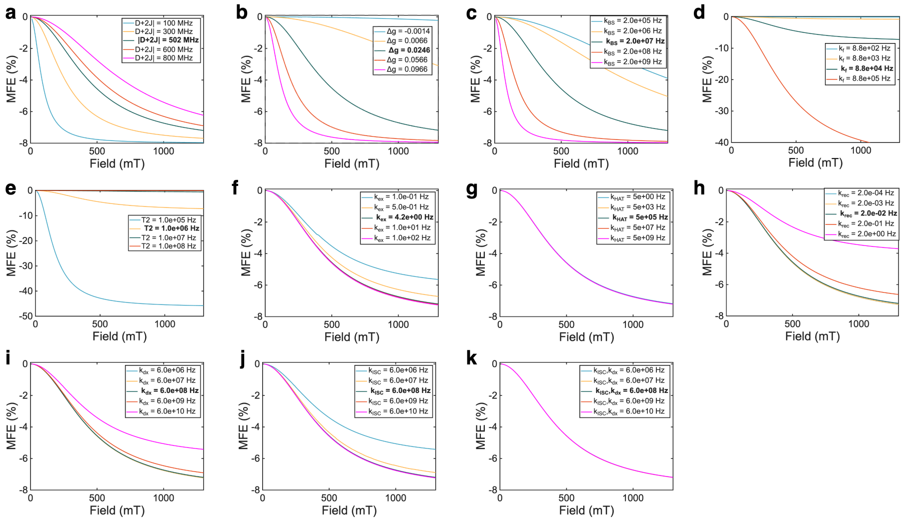


Figure S10: Kinetic MFE simulations varying parameters to show the impact on resulting MFE field dependence. Bolded values in the legend are the values used in the kinetic model fit shown in Fig. 3(d). Magnetic field effects are strongly dependent on the spin parameters ($|D+2J|, \Delta g, T_2$) and the radical pair kinetics ($k_f, k_{BS}$), but are relatively insensitive to the other kinetics of the photocycle unless varied by several orders of magnitude.

#### S.8 Quantum mechanical modeling of radical pair dynamics

Simulations were carried out in Spinach package version 2.13[11,12]. Simulations of spin dynamics evolutions were performed in Liouville space using a spin Hamiltonian for the radical pair that included electron Zeeman interactions of both radicals (with g-values of $g_{FMN,iso}$ and $g_{cyst,iso}$ listed in Table 1) and an isotropic exchange coupling of $J = 250$ MHz. No dipolar tensor was entered, so this calculation is the liquid limit (rapid isotropic tumbling), in which the traceless dipolar tensor averages to zero, so that the full contribution of the $|D + 2J|$ term was accounted for in $J$.

Spin relaxation was described phenomenologically by longitudinal and transverse relaxation terms, and relaxation rates being assumed to be equal for both radical partners in the pair. Kinetics superoperator was constructed assuming the Haberkorn kinetics model[13] with rate constants of evolution from singlet and triplet states equal to $k_{BS}$ and $k_f$, respectively. The time evolution of populations of $S$ and $T_0$ states is shown for varying fields in Fig. S11(a), dependence on longitudinal relaxation rate is shown in Fig. S11(b).

Magnetic field effect was modeled using function rydmr, which was modified to allow for the simulation of triplet-born radical pairs. The singlet yield (the probability of the bound state formation during the whole lifetime of the radical pair) was calculated as[14]:

$$\Phi_S = \int_0^\infty k_{BS} \left\langle \hat{S} \middle| \rho\left(t\right) \right\rangle dt = k_{BS} \mathrm{Im} \left\langle \hat{S} \middle| \hat{\hat{L}}^{-1} \left| \rho_0 \right\rangle \right. \tag{1}$$

where $\hat{S}$ is the singlet projection operator, $\hat{\hat{L}}$ is the Liouvillian superoperator and $\rho_0$ corresponded to the initial state of the radical pair. For the reported simulations the radical pair was assumed to be born in pure $T_0$ state. Under the assumption that the radical pairs in the initial states can equiprobably populate $T_0$ and $T_\pm$, the shape of the magnetic field dependence of the singlet yield does not change its shape, and the singlet yield only changes proportionally (Fig. S11(c)).

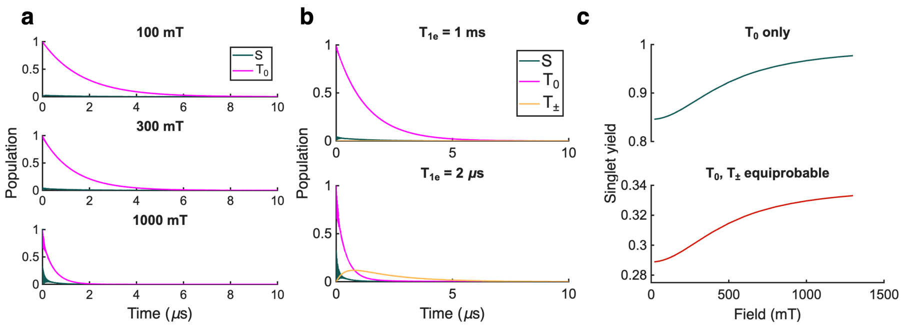


Figure S11: (a, b) Time dependence of singlet ($S$) and triplet ($T_0$ and $T_\pm$) spin state populations during the evolution of the radical pairs at different magnetic fields (a) and at different rates of electron longitudinal relaxation (b). (c) The magnetic field dependence of singlet yield for different assumptions on the initial state of the radical pair: purely $T_0$-born radical pairs (top) and equiprobably-populated $T_0$, $T_+$ and $T_-$ states (bottom).

#### S.9 Optically Detected Magnetic Resonance (ODMR)

Following Abrahams et al.[1], we replicated their MagLOV cw-ODMR spectrum at X-band fields (see Methods for details). Note that our spectrum is significantly broader because of the high microwave power used on our setup. We attempted to do ODMR on AsLOV2, but attempts were unsuccessful (no contrast was observed), likely due to a broad spectrum due to the dipolar and exchange couplings, as well as the highly temperature-dependent fluorescence obscuring any genuine signal. Future work with AsLOV2 cw-ODMR will likely require cryogenic conditions and longer experiment times.

Additionally, cw-ODMR of AsLOV2 would function differently from MagLOV due to the different mechanisms leading to the MFE for each protein. In MagLOV, the presence of a field generally decouples the $T_\pm$ levels from the $S_0$ state, and introducing microwaves recouples the $T_\pm$ levels to the $T_0$ level and thus the $S_0$ state, leading to a recovery of the fluorescence decrease seen in the field on conditions without microwaves. However, in AsLOV2 the $T_\pm$ levels are never coupled to the $S_0$ state in either the field on or field off conditions. When the field is off, the triplet and singlet states are eigenstates of the spin Hamiltonian, as described in the main text, and no interconversion occurs (S.7). When the field is sufficiently high to allow for $T_0 - S_0$ interconversion, the Zeeman energy of the $T_\pm$ states decouples these states from interconversion with $S_0$. While microwaves could couple the $T_\pm$ and $T_0$ states, the effect on the AsLOV2 fluorescence is not clear as this does not recover the initial field off behavior as in the case of MagLOV.

See the Methods section for more details about the attempted experiments.

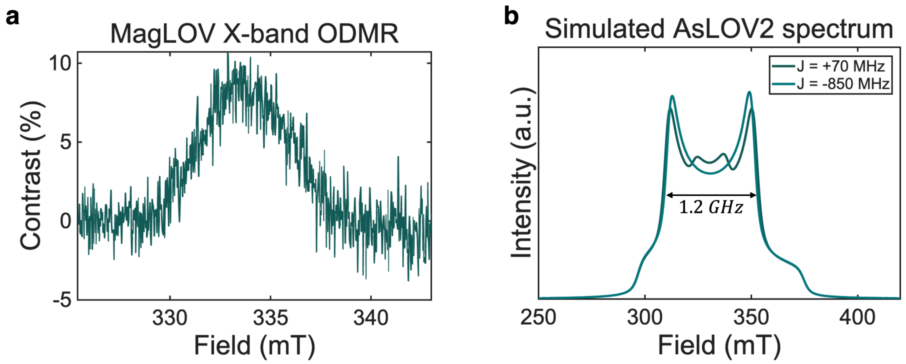


Figure S12: (a) MagLOV ODMR spectrum at X-band. A background function was fit and subtracted from the spectrum to account for drift in fluorescence from sample heating during the experiment. (b) Simulated AsLOV2 radical pair spectra for $d = 728$ MHz with $J = +70$ MHz and $J = -850$ MHz. Both spectra show a linewidth of 1.2 GHz, so that efficient saturation of the AsLOV2 radical pair is not possible with our X-band system.

#### S.10 Electronic structure of dark and lit states

The magnetic field effect (MFE) reported in the main text requires a photogenerated, spin-correlated FMN–cysteine radical pair (SCRP) on the photoadduct pathway, formed by hydrogen atom transfer from C450 to the photoexcited flavin. The MFE itself is diagnostic of this radical pair, but the strong dipolar and exchange coupling that set the high-field onset also preclude its direct observation by EPR/ODMR (main text and S.9). To provide an independent, structure-based check on the two photochemical premises that underlie the kinetic model of Fig. 3(c) - (i) that the dark and lit states carry the distinct UV–Vis signatures used as the MFE readout in Fig. 2, and (ii) that low-lying flavin→cysteine charge-transfer (CT) states are available to seed the electron-transfer step, we carried out time-dependent density functional theory (TDDFT) on a truncated FMN–cysteine dyad (details in Methods M.11).

*Assignment of the probe bands.* With a single shift of $-0.375$ eV the calculated spectra reproduce both states (Fig. 2(a,b); Fig. S13).

**Dark state, 447 nm.** The band is a single transition ($S_1$, $f = 0.44$; MO 142/143 → LUMO 147, weights 0.61/0.30). Its hole is 93% on the isoalloxazine and its particle is the flavin $\pi^*$ orbital with large N5 and C4a coefficients (Fig. S13). It is a local flavin $\pi\pi^*$ excitation, which requires conjugation through C4a.

**Lit state, 310 nm.** In the adduct C4a is $sp^3$ and the 447 nm transition is gone. The absorbance near 310 nm comes from $S_5$ (291 nm, $f = 0.17$; about 60% of the calculated 310 nm absorbance) and $S_3$ (338 nm, $f = 0.21$; about 40%), both into the uracil-type $\pi^*$ of the reduced flavin (Fig. S13). $S_3$ has substantial sulfur character in its hole (cysteine 41%). It lies at the same wavelength as a

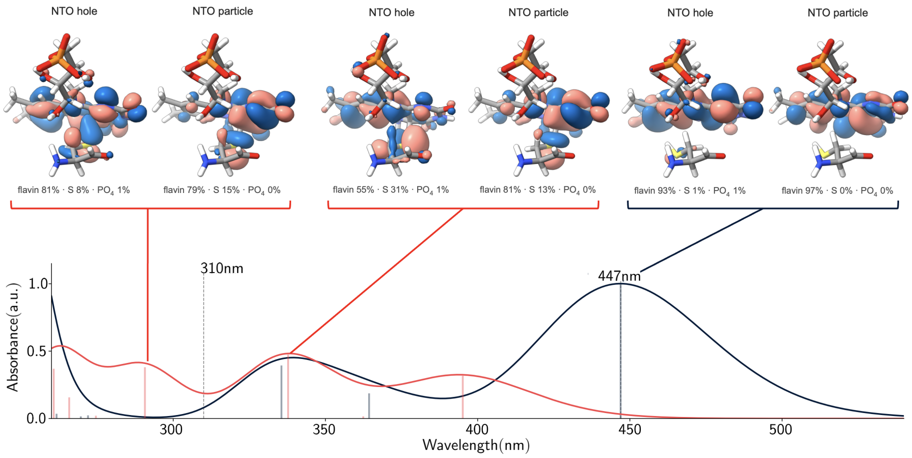


Figure S13: Transitions behind the 447 nm (dark) and 310 nm (lit) probe bands (TDA $\omega$B97X-D4/def2-TZVP, CPCM $\varepsilon = 4$; shifted by $-0.375$ eV).

weaker dark-state transition ($S_8$, 336 nm), so the dark/lit contrast at 310 nm comes mainly from $S_5$. $S_1$ of the adduct (395 nm, $f = 0.14$) corresponds to the measured 380 nm band.

The calculated lit$-$dark difference therefore has a strong bleach at 447 nm and gains near 380 nm and below 340 nm, as measured (Fig. 2(a)).

*Evolution along the path.* Between 4.35 and 3.65 Å (which includes the 4.39 and 3.75 Å rotamers of 2V1A) the lowest bright band stays at 445–452 nm, and the 420–500 nm absorbance falls by less than 20% (Fig. 2(c)). As the thiol proton approaches N5 (3.5–3.3 Å) the band moves to 460–490 nm. Between 3.1 and 2.4 Å the closed-shell description fails (below), and the spectra in this region are qualitative. The adduct bands at 380 and 310 nm appear only at 2.1 Å and below, where 14% of the dark 420–500 nm absorbance remains. Bond formation, proton transfer and C4a pyramidalization change together along the interpolated path, so the spectral switch marks loss of conjugation at C4a, not a property of the S–C4a distance alone.
